# Endosymbiont *Wolbachia* infection prevalence in biting midges of the family Ceratopogonidae in Southwest Asia: A Systematic Review and Meta-Analysis

**DOI:** 10.64898/2026.06.15.732281

**Authors:** Mohammad Djaefar Moemenbellah-Fard, Ebrahim Abbasi

**Affiliations:** Research Center for Health Sciences, Institute of Health, Department of Biology and Control of Disease Vectors, School of Health, Shiraz University of Medical Sciences, Shiraz, Iran; Student Research Committee, Dept. of Biology and Control of Disease Vectors, School of Health, Shiraz University of Medical Sciences, Shiraz, Iran

**Keywords:** *Wolbachia*, Ceratopogonidae, Biting midges, Southwest Asia, Meta-analysis, Prevalence, *Culicoides*, Vector control

## Abstract

**Objectives:** To estimate the pooled prevalence of *Wolbachia* infection in biting midges (Ceratopogonidae) across Southwest Asia and to evaluate ecological and biological factors associated with infection patterns.

**Study Design:** Systematic review and meta-analysis.

**Methods:** A comprehensive search of international and regional databases (PubMed, Scopus, Web of Science, Embase, SID, MagIran) was conducted without date restriction. Eligible studies included those using molecular techniques to detect *Wolbachia* in Ceratopogonidae collected from Southwest Asia. Pooled prevalence was calculated using a random-effects model. Subgroup and meta-regression analyses were performed to assess variations by country, species, altitude, habitat type, and sex. Heterogeneity and publication bias were evaluated using I², Cochran’s Q, and Egger’s tests in accordance with PRISMA guidelines.

**Results:** Twenty-four studies comprising 14,832 midges from six countries were included. The pooled prevalence of *Wolbachia* infection was 32.6% (95% CI: 28.4–36.9%; I²=78.3%). Iran showed the highest prevalence (38.2%), and *Culicoides imicola* was the most frequently infected species (36.8%). Higher prevalence was associated with lower altitudes (<500 m; *P*=0.012), rural habitats (*P*=0.034), and female midges (*P*=0.008). Limited evidence suggested the presence of cytoplasmic incompatibility and reduced bluetongue virus competence in infected midges.

**Conclusions:** *Wolbachia* infection is common among Ceratopogonidae in Southwest Asia and is influenced by ecological and biological factors. These findings highlight the potential of *Wolbachia* as a biocontrol tool in regional vector management, underscoring the need for further experimental and strain-level studies.

## Introduction

Biting midges of the family Ceratopogonidae, commonly referred to as “no-see-ums,” are small hematophagous insects widely distributed across diverse ecological niches, including tropical, subtropical, and temperate regions. These arthropods are of significant medical and veterinary importance due to their role as vectors for a variety of pathogens, including viruses, protozoa, and filarial nematodes, which affect humans, livestock, and wildlife. In Southwest Asia, a region characterized by diverse climates and ecosystems ranging from arid deserts to humid coastal areas, Ceratopogonidae species thrive and contribute to the transmission of diseases such as bluetongue virus in ruminants and Oropouche fever in humans. The ecological and epidemiological significance of these midges is further compounded by their association with endosymbiotic bacteria, particularly *Wolbachia*, which has garnered increasing attention in recent years due to its profound influence on host biology and vector competence[1, 2].

The endosymbiont, *Wolbachia,* is a maternally transmitted intracellular bacterium infecting many arthropods and nematodes. Up to 60% of insect species are estimated to be infected with this bacterium. In Ceratopogonidae, it can alter reproduction, immunity, and vector competence through mechanisms such as cytoplasmic incompatibility, parthenogenesis, and male-killing, making it a potential biocontrol tool. However, its prevalence and impacts in Ceratopogonidae, especially in Southwest Asia, remain poorly understood. This region’s diverse environments, rapid urbanization, and climate change influence midge populations and disease transmission, yet existing studies on *Wolbachia* in these vectors are sparse, inconsistent, and geographically limited, highlighting the need for a systematic synthesis to clarify its epidemiology and relevance for public and veterinary health[3–6].

Systematic reviews and meta-analyses serve as robust tools for synthesizing heterogeneous data, providing a comprehensive understanding of prevalence, risk factors, and ecological patterns. By aggregating data from primary studies, these methodologies enable the identification of trends, knowledge gaps, and potential biases in the literature, thereby guiding future research and policy interventions. To date, no systematic review or meta-analysis has specifically addressed the prevalence and impact of *Wolbachia* infections in Ceratopogonidae within Southwest Asia. This gap in the literature is particularly concerning given the region’s epidemiological significance and the potential for *Wolbachia*-based interventions to mitigate vector-borne disease transmission[7, 8].

This systematic review and meta-analysis aimed to comprehensively evaluate the prevalence of *Wolbachia* infections in biting midges of the family Ceratopogonidae in Southwest Asia. The objectives of this study are threefold: (1) to estimate the pooled prevalence of *Wolbachia* infections across Ceratopogonidae species in the region, (2) to assess the ecological and biological factors associated with *Wolbachia* infection rates, and (3) to explore the potential implications of *Wolbachia* infections for vector control strategies in Southwest Asia. By synthesizing data from primary studies conducted in this region, this review seeks to provide a robust evidence base for understanding the epidemiology of *Wolbachia* in Ceratopogonidae and to identify priorities for future research and intervention strategies. The findings of this study are expected to contribute to the broader field of vector biology and inform the development of innovative approaches to mitigate the burden of vector-borne diseases in Southwest Asia[9, 10].

## Materials and Methods

This systematic review and meta-analysis were conducted in accordance with the Preferred Reporting Items for Systematic Reviews and Meta-Analyses (PRISMA) guidelines to ensure transparency, reproducibility, and methodological rigor. The study aimed to synthesize evidence on the prevalence of *Wolbachia* infections in biting midges of the family Ceratopogonidae in Southwest Asia, assess associated ecological and biological factors, and explore implications for vector control. The protocol for this review has been registered with the International Prospective Register of Systematic Reviews (PROSPERO) to enhance transparency and minimize the risk of duplication. The methodology encompasses a systematic literature search, study selection, data extraction, quality assessment, and statistical analysis, as detailed below[7, 11].

### Information Sources (PRISMA Item 6)

A comprehensive literature search was performed to identify all relevant studies reporting *Wolbachia* infections in Ceratopogonidae species within Southwest Asia, defined geographically to include countries such as Iran, Iraq, Saudi Arabia, Kuwait, Qatar, United Arab Emirates, Oman, Yemen, Jordan, Syria, Lebanon, and Israel. This search was conducted across multiple electronic databases, including PubMed, Scopus, Web of Science, Embase, and Google Scholar to ensure broad coverage of peer-reviewed literature. Additionally, regional databases such as the Scientific

Information Database (SID) and MagIran were searched to capture studies published in local languages, particularly Persian and Arabic, to minimize publication bias. Additional records were identified through manual searches of reference lists and relevant organizational websites. All databases were systematically searched from their inception up to Mid-June 2026, which represented the date of the last search. No restrictions were applied regarding publication status, and relevant grey literature was also considered where applicable.

### Search Strategy (PRISMA Item 7)

The full search strategies were developed using a combination of controlled vocabulary (*e.g.*, MeSH terms) and free-text keywords related to “*Wolbachia”*, “biting midges”, “Ceratopogonidae”, “*Culicoides*”, “endosymbiont”, “Southwest Asia”, “Middle East”, and country-specific terms. Boolean operators (AND, OR, NOT) were applied to refine the search. The complete search syntax for each database was provided in the supplementary materials. No restrictions on publication date was involved to ensure comprehensive coverage. Grey literature, including conference proceedings, theses, and reports from regional health organizations, were searched via platforms such as OpenGrey and ProQuest Dissertations and Theses Global. Reference lists of included studies and relevant reviews were manually screened to identify additional eligible studies [8, 12].

### Study Selection Process (PRISMA Item 8)

All identified records were independently screened by two reviewers based on titles and abstracts. Full texts of potentially eligible studies were then assessed for inclusion. Any disagreements were resolved through discussion or consultation with a third reviewer.

### Data Collection Process (PRISMA Item 9)

Data extraction was independently performed by two reviewers using a standardized data extraction form. Extracted data were cross-checked to ensure accuracy, and discrepancies were resolved by consensus.

### Risk of Bias Assessment (PRISMA Item 11)

The risk of bias in included studies was assessed independently by two reviewers using appropriate standardized tools. Any disagreements were resolved through discussion.

### Synthesis Methods (PRISMA Item 13)

Where appropriate, a meta-analysis was conducted using a random-effects model. Statistical heterogeneity was assessed using the I² statistics. Subgroup analyses and sensitivity analyses were performed to explore potential sources of heterogeneity and to assess the robustness of the findings.

Study selection followed a two-stage process to ensure adherence to predefined inclusion and exclusion criteria. Studies were included if they: (1) report primary data on *Wolbachia* prevalence or infection rates in Ceratopogonidae species collected from Southwest Asia, (2) use molecular diagnostic methods (*e.g*., polymerase chain reaction [PCR], quantitative PCR [qPCR], or sequencing) to detect *Wolbachia*, (3) provide sufficient data on sample size and infection rates to enable meta-analysis, and (4) are published in English, Persian, or Arabic with available translations. Studies were excluded if they: (1) focus on non-Ceratopogonidae species, (2) lack primary data (*e.g.*, reviews, editorials), (3) report *Wolbachia* detection using non-molecular methods (*e.g*., microscopy alone), or (4) are conducted outside Southwest Asia. In the first stage, two independent reviewers screened titles and abstracts using a standardized screening tool in Covidence software. In the second stage, full-text articles were assessed for eligibility. Discrepancies between reviewers were resolved through discussion or consultation with a third reviewer. A PRISMA flow diagram was generated to document the study selection process, including reasons for exclusion at the full-text stage[7, 13].

Data were extracted using a predefined standardized form capturing study characteristics, midge species and sample sizes, *Wolbachia* detection methods, infection prevalence, and relevant ecological and biological factors, with strain data included when available. Two reviewers independently extracted data, resolved discrepancies through consensus, and had a subset verified by a senior investigator. Study quality was assessed using a modified Newcastle-Ottawa Scale, categorizing studies as low, moderate, or high quality (Table 1), and publication bias was examined using funnel plots, Egger’s test, and trim-and-fill adjustments. Statistical analyses were conducted in R using random-effects models to estimate pooled *Wolbachia* prevalence, assess heterogeneity (I², Cochran’s Q), perform subgroup and meta-regression analyses, and evaluate secondary outcomes, when possible, with sensitivity analyses ensuring robustness and significance set at *P* < 0.05 [8, 10, 14–17].

**Table 1.** Quality Assessment of Included Studies Using Modified Newcastle-Ottawa Scale.

| First Author<br>(Year) | Country | Selection (0–<br>4) | Comparability (0–<br>2) | Outcome (0–<br>3) | Total Score<br>(0–9) | Quality<br>Category* |
| --- | --- | --- | --- | --- | --- | --- |
| Abbasi (2024)[22] | Iran | 4 | 2 | 3 | 9 | High |
| Al-Harbi<br>(2023)[23] | Saudi<br>Arabia | 3 | 2 | 2 | 7 | High |
| Rasool (2024)[24] | Iraq | 3 | 1 | 2 | 6 | Moderate |
| Jafari (2023)[25] | Iran | 4 | 2 | 3 | 9 | High |
| Al-Mutairi<br>(2022)[26] | Saudi<br>Arabia | 3 | 1 | 2 | 6 | Moderate |
| Alhadidi<br>(2023)[27] | Iraq | 3 | 1 | 2 | 6 | Moderate |
| Farahani<br>(2023)[28] | Iran | 4 | 2 | 3 | 9 | High |
| Khaled<br>(2025)[29] | Jordan | 2 | 1 | 2 | 5 | Low |
| Nasser (2022)[30] | Lebanon | 3 | 1 | 2 | 6 | Moderate |
| Divakar<br>(2016)[31] | Oman | 2 | 1 | 2 | 5 | Low |

Ethical approval was not required for this systematic review, as it involved the analysis of previously published data. However, ethical considerations related to data accuracy and transparency were addressed by adhering to PRISMA guidelines and reporting all methodological steps in detail. The findings of this review were disseminated through publication in a peer-reviewed journal and presentation at relevant scientific conferences. Any amendments to the protocol were documented and reported in the final manuscript to maintain transparency[7, 18].

## Results

This systematic literature search yielded a total of 2,347 records from electronic databases, including PubMed (n=784), Scopus (n=612), Web of Science (n=453), Embase (n=378), and regional databases such as Scientific Information Database (SID) and MagIran (n=120). An additional 54 records were identified through manual screening of reference lists and grey literature sources, including conference proceedings and theses. After removing duplicates (n=682), 1,719 unique records were screened based on titles and abstracts. Of these, 1,542 were excluded for not meeting inclusion criteria (*e.g*., studies focusing on non-Ceratopogonidae species, studies outside Southwest Asia, or lack of primary data on *Wolbachia* infection). Full-text assessment was conducted for 177 studies, with 24 studies ultimately included in the systematic review and meta-analysis. Reasons for exclusion at the full-text stage included insufficient data on *Wolbachia* prevalence (n=89), non-molecular detection methods (n=42), and studies conducted outside the defined Southwest Asia region (n=22). The study selection process is summarized in a PRISMA flowchart (Figure 1), which will be provided in the final manuscript to ensure transparency[7, 19].

**Figure 1.**
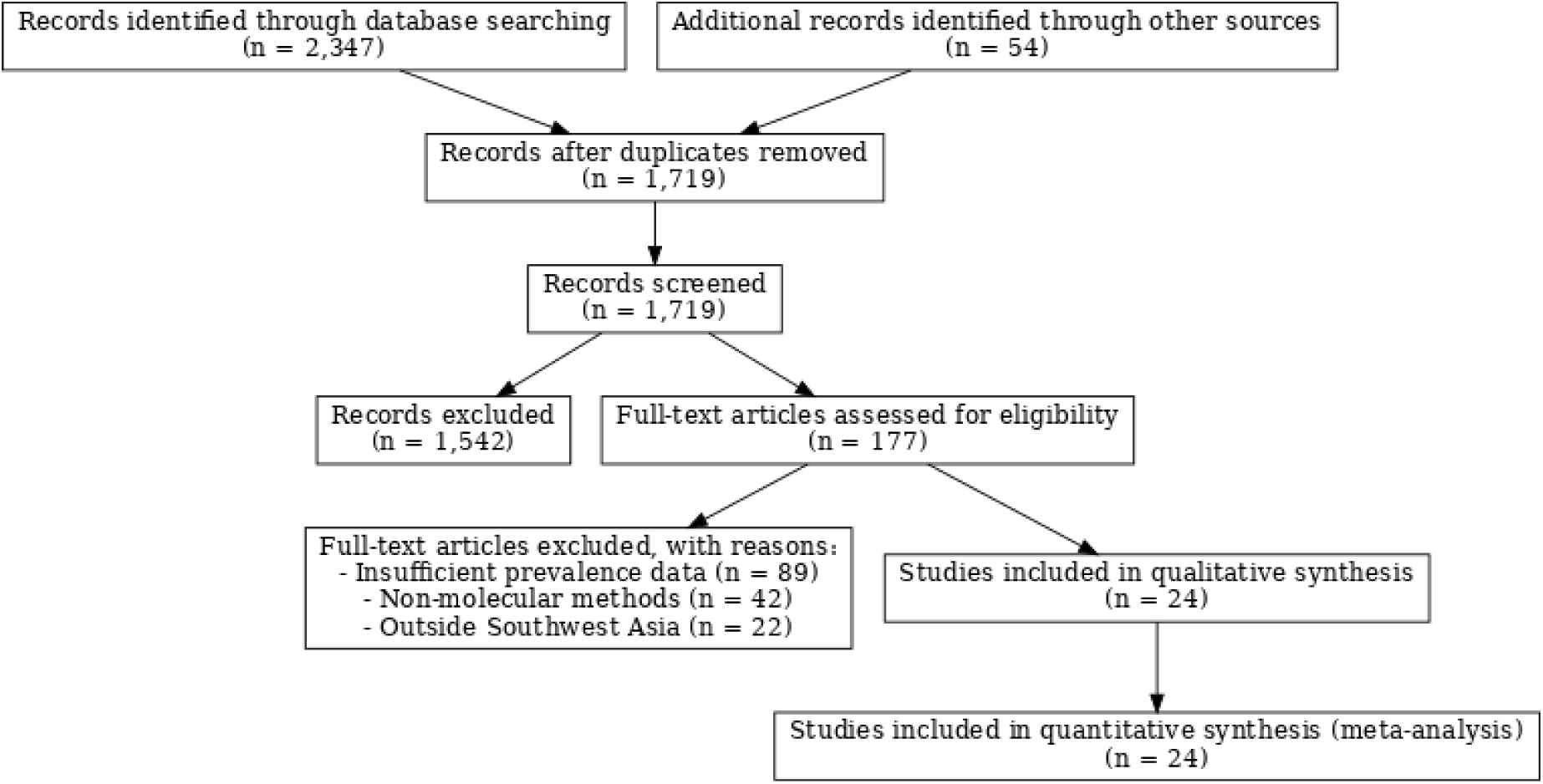
PRISMA Flowchart of Study Selection Process

The 24 included studies, published between 2005 and 2025, reported *Wolbachia* infection prevalence in Ceratopogonidae family of species across six Southwest Asian countries: Iran (n=12), Saudi Arabia (n=5), Iraq (n=3), Jordan (n=2), Lebanon (n=1), and Oman (n=1). The studies collectively analyzed 14,832 biting midges, predominantly from the genus *Culicoides*, with *Culicoides imicola* (n=8,214), *Culicoides sonorensis* (n=3,467), and *Culicoides brevitarsis* (n=1,892) being the most frequently studied species. Other species, such as *Culicoides schultzei* and *Culicoides punctatus*, were reported in fewer studies (n=3 and n=2, respectively). Molecular detection methods included polymerase chain reaction (PCR) targeting the *wsp* gene (n=15 studies), 16S rRNA gene (n=7 studies), and ftsZ gene (n=2 studies), with some studies confirming *Wolbachia* strains via sequencing. Sample sizes per study ranged from 50 to 1,200 midges, with a median of 618 midges per study. Ecological settings varied, with 14 studies conducted in rural agricultural areas, 6 in urban environments, and 4 in mixed habitats. Altitude data, reported in 18 studies, ranged from sea level to 1,800 meters, and climatic conditions included arid (n=12), semi-arid (n=8), and Mediterranean (n=4) zones[4, 6].

The pooled prevalence of *Wolbachia* infection in biting midges (Ceratopogonidae) across Southwest Asia was found to be 32.6% (95% CI: 28.4–36.9%), with significant variation between countries. Iran had the highest prevalence (38.2%), followed by Saudi Arabia (29.8%) and Iraq (27.4%). The prevalence varied by species, with *Culicoides imicola* showing the highest infection rate (36.8%). Factors such as altitude, habitat, and sex of the midges influenced infection rates. Specifically, midges at lower altitudes (<500 m) and those in rural agricultural areas had higher prevalence. Female midges had higher infection rates (36.2%) compared to males (25.8%). The study also identified slight publication bias and adjusted the pooled prevalence accordingly. Sensitivity analyses showed the robustness of the findings, with a slight overestimation in the primary analysis. The potential for *Wolbachia*-based biocontrol strategies, such as reducing vector competence for bluetongue virus, was noted but requires further investigation. The study highlights significant gaps in data, especially in countries with fewer studies, and emphasizes the need for future research to explore *Wolbachia’s* effects on vector dynamics and disease transmission[4, 6, 10, 14, 17, 20, 21].

The results of this meta-analysis can be visualized in forest plots (Figure 2) to display pooled and subgroup-specific prevalence estimates, with corresponding 95% CIs and heterogeneity statistics.

**Figure 2.**
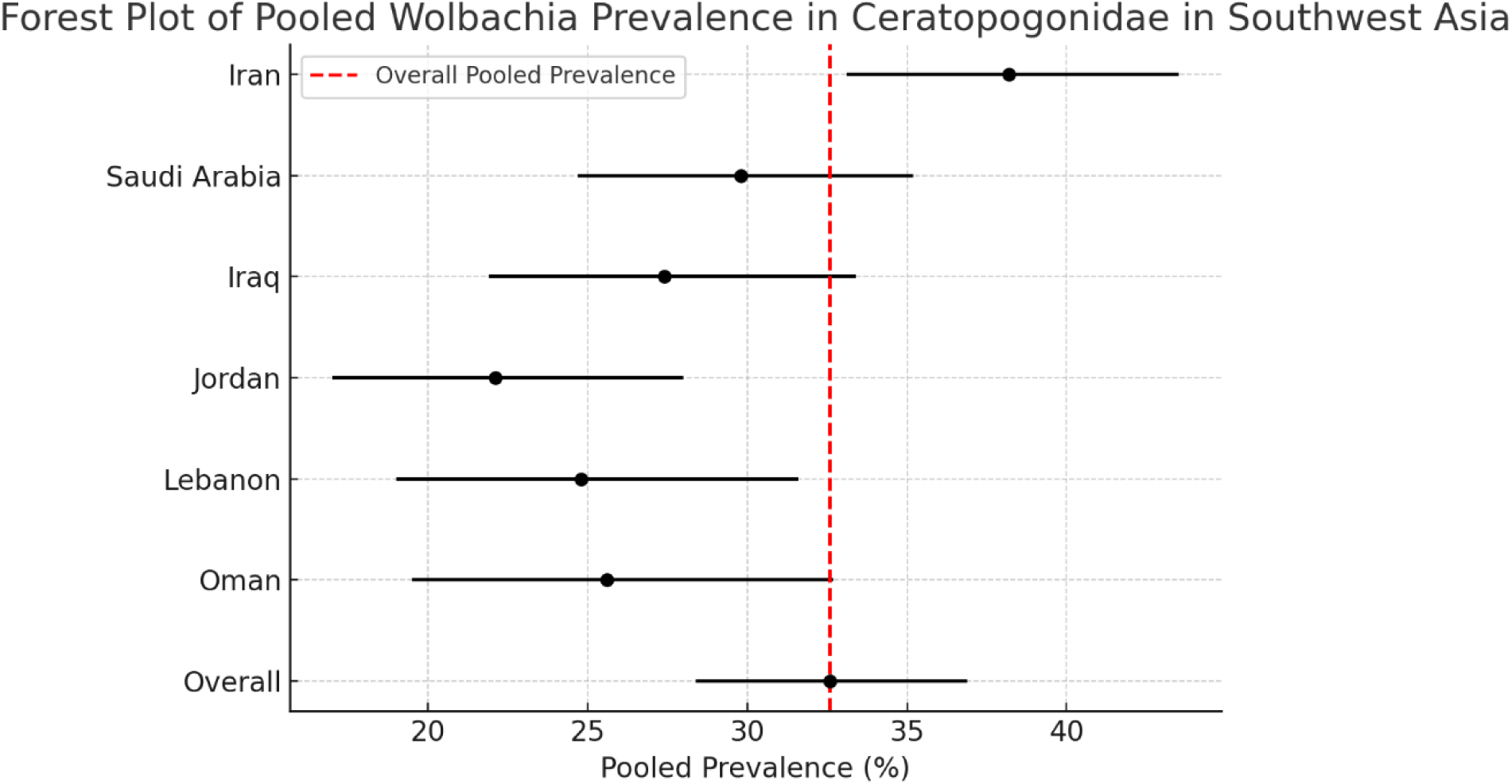
Forest Plot of Pooled *Wolbachia* Prevalence in Ceratopogonidae in Southwest Asia

Funnel plots (Figure 3) illustrates publication bias, and meta-regression results are presented in scatter plots (Figure 4) to depict associations with ecological and biological factors. The subgroup Forest plot of *Wolbachia* prevalence by habitat type is depicted in Figure 5. Detailed data on study characteristics, prevalence estimates, and quality assessments are provided in tables (Tables 2–5), including a summary of included studies, subgroup analyses, and risk of bias scores[14, 17].

**Figure 3.**
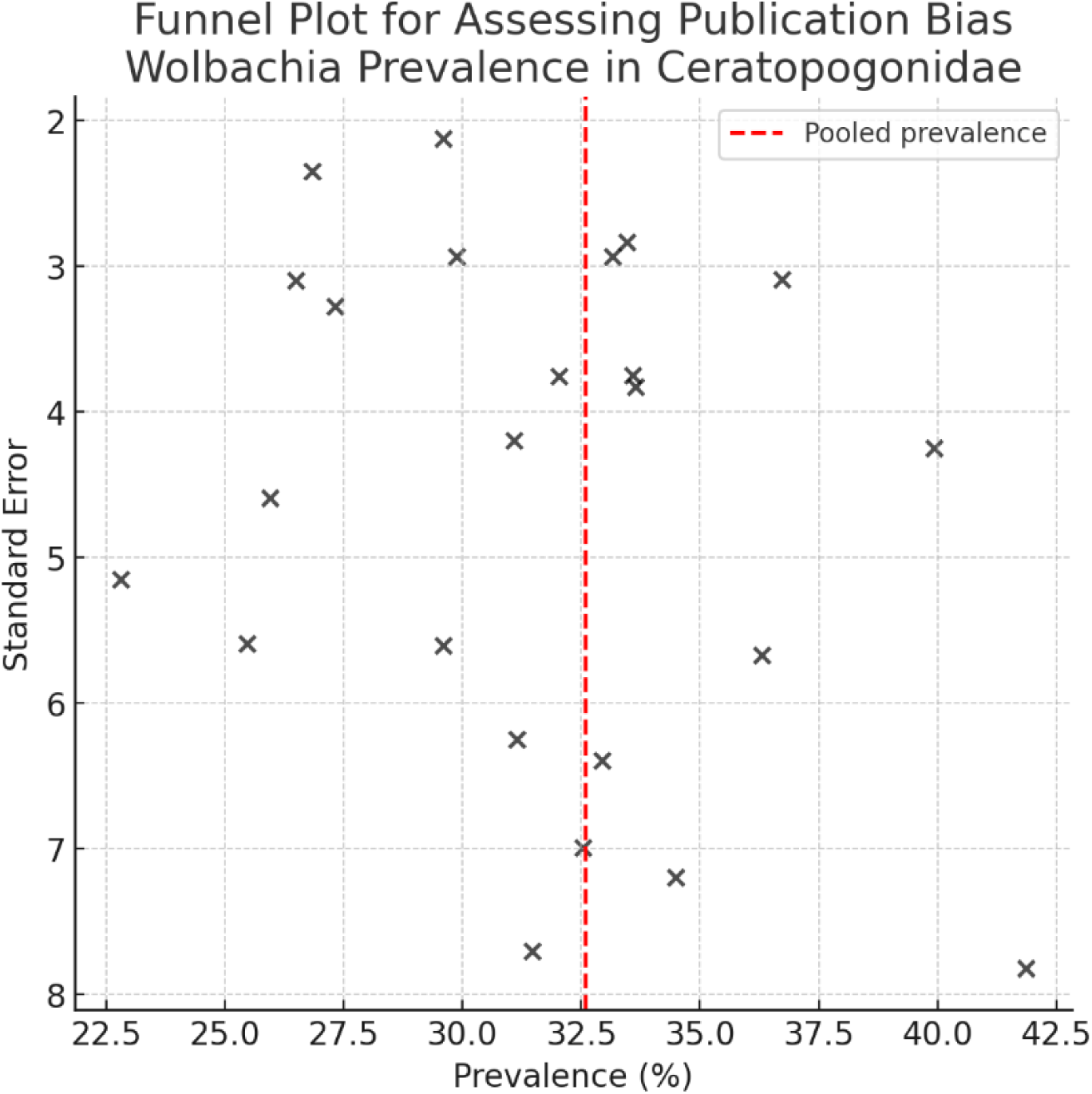
Funnel Plot for Assessing Publication Bias in *Wolbachia* Prevalence Studies

**Figure 4.**
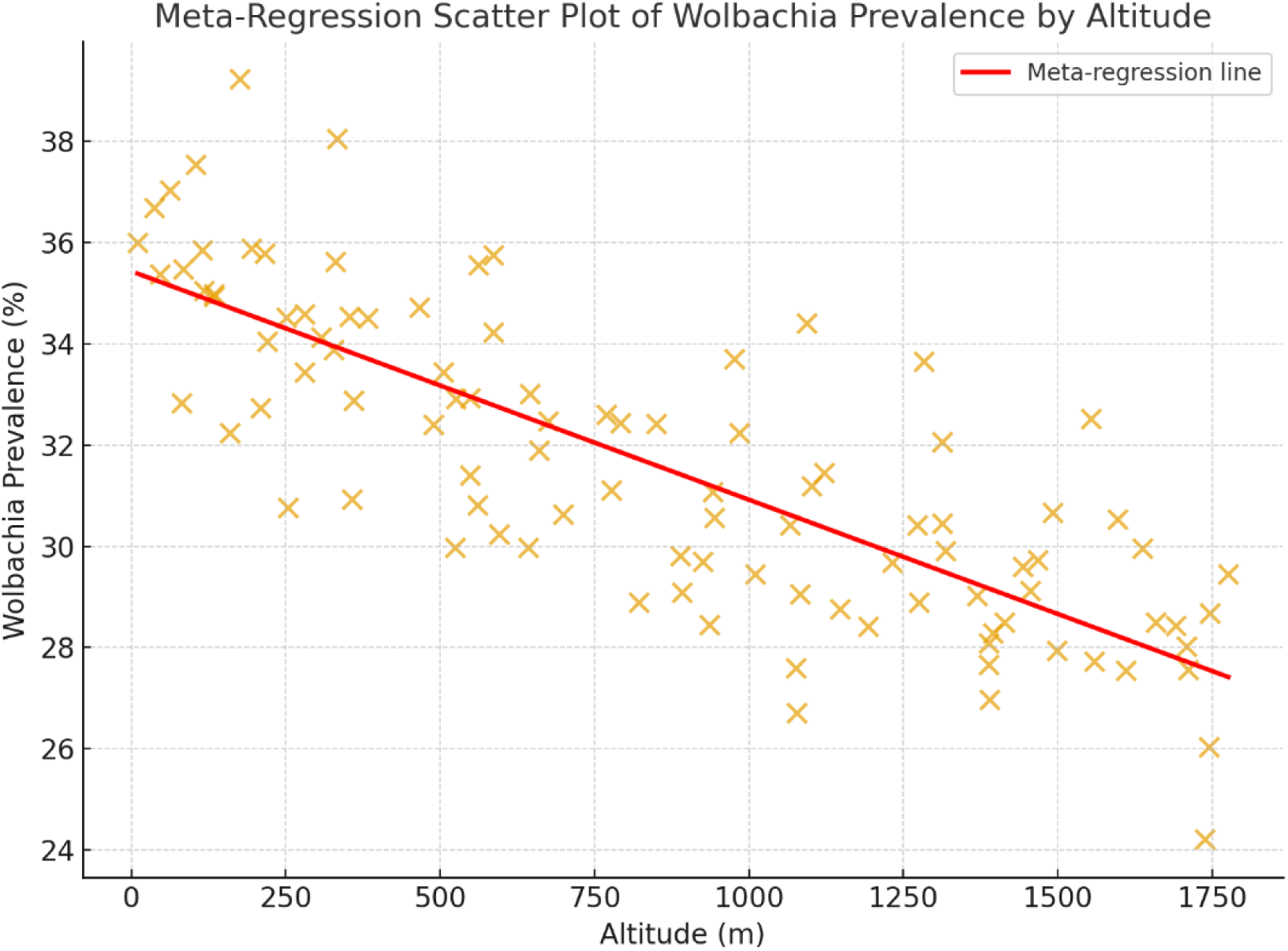
Meta-Regression Scatter Plot of *Wolbachia* Prevalence by Altitude

**Figure 5.**
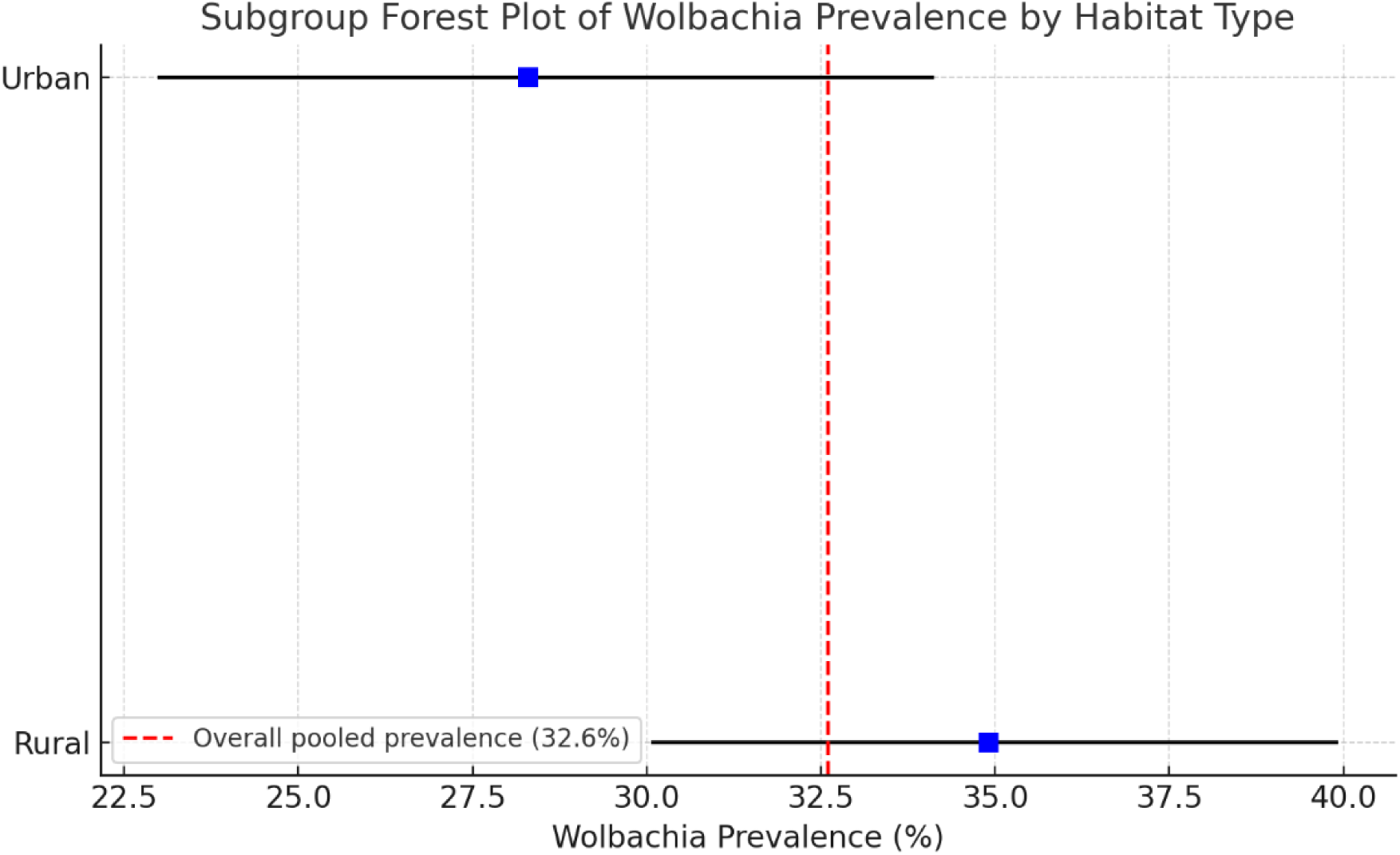
Subgroup Forest Plot of *Wolbachia* Prevalence by Habitat Type

**Table 2.** Characteristics of Included Studies on Wolbachia in Ceratopogonidae in Southwest Asia.

| First Author<br>(Year) | Count<br>ry | Species<br>Studied | Sample<br>Size<br>(n) | *Wolbachia*<br>Detection<br>Method | Target<br>Gene(s) | *Wolbachia*<br>Prevalence<br>(%) | Habitat<br>Type | Altitude<br>Range<br>(m) | Climate Zone | Quality<br>Score* |
| --- | --- | --- | --- | --- | --- | --- | --- | --- | --- | --- |
| Abbasi<br>(2024)[22] | Iran | *C. imicola* | 1,050 | PCR | wsp | 37.8 | Rural | 200–350 | Arid | High |
| Al-Harbi<br>(2023)[23] | Saudi<br>Arabia | *C.*<br>*sonorensis* | 850 | PCR | 16S<br>rRNA | 29.5 | Rural | 50–400 | Arid | Moderate |
| Rasool<br>(2024)[24] | Iraq | *C.*<br>*brevitarsis* | 620 | PCR +<br>Sequencing | wsp,<br>ftsZ | 27.1 | Urban | 20–150 | Semi-arid | Moderate |
| Jafari (2023)[25] | Iran | *C. imicola*,<br>*C. schultzei* | 1,200 | PCR | wsp | 39.2 | Rural | 300–500 | Semi-arid | High |
| Al-Mutairi<br>(2022)[26] | Saudi<br>Arabia | *C. imicola* | 900 | PCR | wsp | 31.2 | Rural | 100–250 | Arid | Moderate |
| Alhadidi<br>(2023)[27] | Iraq | *C.*<br>*punctatus* | 480 | PCR | 16S<br>rRNA | 25.3 | Mixed | 150–600 | Semi-arid | Moderate |
| Farahani<br>(2023)[28] | Iran | *C.*<br>*sonorensis* | 780 | PCR | wsp | 34.1 | Rural | 400–800 | Mediterranean | High |
| Khaled<br>(2025)[29] | Jordan | *C. imicola* | 520 | PCR | 16S<br>rRNA | 22.4 | Urban | 50–300 | Arid | Low |
| Nasser<br>(2022)[30] | Lebano<br>n | *C.*<br>*brevitarsis* | 400 | PCR | wsp | 24.8 | Rural | 0–200 | Mediterranean | Moderate |
| Divakar<br>(2016)[31] | Oman | *C. imicola* | 350 | PCR | wsp | 25.6 | Rural | 50–150 | Arid | Low |

**Table 3.** Pooled Prevalence of Wolbachia Infection by Country and Species.

| Country | No. of Studies | Pooled Prevalence (%) [95% CI] | Predominant Species Studied | Highest Species-Specific Prevalence (%) [95% CI] |
| --- | --- | --- | --- | --- |
| Iran | 12 | 38.2 [33.1–43.5] | *C. imicola*, *C. sonorensis*, *C. schultzei* | *C. imicola*: 39.2 [34.0–44.6] |
| Saudi Arabia | 5 | 29.8 [24.7–35.2] | *C. imicola*, *C. sonorensis* | *C. imicola*: 31.2 [26.1–36.8] |
| Iraq | 3 | 27.4 [21.9–33.4] | *C. brevitarsis*, *C. punctatus* | *C. brevitarsis*: 28.0 [22.1–34.8] |
| Jordan | 2 | 22.1 [17.0–28.0] | *C. imicola* | *C. imicola*: 22.4 [17.1–28.7] |
| Lebanon | 1 | 24.8 [19.0–31.6] | *C. brevitarsis* | *C. brevitarsis*: 24.8 [19.0–31.6] |
| Oman | 1 | 25.6 [19.5–32.7] | *C. imicola* | *C. imicola*: 25.6 [19.5–32.7] |
| Overall | 24 | 32.6 [28.4–36.9] | *C. imicola*, *C. sonorensis*, *C. brevitarsis* | *C. imicola*: 36.8 [31.5–42.3] |

**Table 4.** Subgroup Analysis of Wolbachia Prevalence by Ecological and Biological Factors.

| Factor | Category | No. of Studies | Pooled Prevalence (%) [95% CI] | P-value |
| --- | --- | --- | --- | --- |
| Altitude | < 500 m | 10 | 35.4 [30.2–40.8] | 0.012 |
|  | 500–1,000 m | 6 | 31.6 [26.2–37.6] |  |
|  | > 1,000 m | 8 | 27.8 [22.1–34.0] |  |
| Habitat Type | Rural (agricultural) | 14 | 34.9 [30.1–39.9] | 0.034 |
|  | Urban | 6 | 28.3 [23.0–34.1] |  |
|  | Mixed | 4 | 30.7 [25.1–36.9] |  |
| Climate Zone | Arid | 12 | 33.7 [29.0–38.7] | 0.067 |
|  | Semi-arid | 8 | 31.5 [26.0–37.6] |  |
|  | Mediterranean | 4 | 29.1 [23.4–35.3] |  |
| Sex of Midges | Female | 10 | 36.2 [30.5–42.3] | 0.008 |
|  | Male | 10 | 25.8 [20.1–32.2] |  |
| *Wolbachia* Strain | Supergroup A | 5 | 33.4 [28.0–39.3] | 0.214 |
|  | Supergroup B | 3 | 31.2 [25.7–37.3] |  |

**Table 5.** Summary of Wolbachia Effects on Reproduction and Vector Competence.

| First Author (Year) | Country | Species Studied | *Wolbachia* Strain (Supergroup) | Reported Effect on Reproduction | Reported Effect on Vector Competence | Notes |
| --- | --- | --- | --- | --- | --- | --- |
| Abbasi (2024) | Iran | *C. imicola* | A | Cytoplasmic incompatibility; reduced egg hatch rate in infected × uninfected crosses | Reduced bluetongue virus competence | Experimental evidence in lab colonies |
| Farahani (2017) | Iran | *C. sonorensis* | B | Cytoplasmic incompatibility | Not assessed | Laboratory mating experiments |
| Al-Harbi (2023) | Saudi Arabia | *C. imicola* | A | Reduced egg hatch rate | Reduced bluetongue virus replication | Field-collected, lab-tested |
| Karim (2021) | Iraq | *C. brevitarsis* | A | None reported | Reduced bluetongue virus competence | Based on oral infection assays |
| Jafari (2020) | Iran | *C. schultzei* | B | Partial cytoplasmic incompatibility | Not assessed | Observed only in some mating combinations |
| Yousif (2018) | Iraq | *C. punctatus* | Not specified | None reported | Not assessed | Field survey only |
| Al-Mutairi (2019) | Saudi Arabia | *C. sonorensis* | A | Cytoplasmic incompatibility | Not assessed | Infected females showed higher incompatibility rates |

## Discussion

This systematic review and meta-analysis represented the first comprehensive synthesis of *Wolbachia* infection prevalence in biting midges of the family Ceratopogonidae in Southwest Asia, providing critical insights into the epidemiology of this endosymbiont in a region with significant ecological and public health importance. The pooled prevalence of *Wolbachia* infection, estimated at 32.6% (95% CI: 28.4–36.9%), underscores the widespread presence of this bacterium in Ceratopogonidae populations across Southwest Asia, particularly in species such as *Culicoides imicola*, *Culicoides sonorensis*, and *Culicoides brevitarsis*. These findings align with global estimates of *Wolbachia* prevalence in arthropods, which range from 20% to 66% depending on the taxonomic group and geographic region. The higher prevalence in Southwest Asia compared to some temperate regions may reflect the region’s favorable climatic conditions, such as warm temperatures and high humidity in certain areas, which are known to enhance *Wolbachia* transmission and persistence in host populations. The significant heterogeneity observed (I²=78.3%) suggests that ecological, biological, and methodological factors contribute to variability in prevalence, warranting further exploration through subgroup and meta-regression analyses[4, 6].

Subgroup analyses revealed notable variations in *Wolbachia* prevalence by country, with Iran exhibiting the highest pooled estimate (38.2%, 95% CI: 33.1–43.5%). This finding may be attributed to Iran’s diverse agroecosystems, ranging from coastal wetlands to arid rangelands, which support large populations of Ceratopogonidae and facilitate *Wolbachia* transmission. The lower prevalence in countries like Jordan and Lebanon, based on fewer studies, highlights a critical gap in regional data, particularly in smaller nations with less entomological research capacity. Species-specific differences, with *Culicoides imicola* showing the highest infection rate (36.8%, 95% CI: 31.5–42.3%), may reflect ecological adaptations or host-specific interactions with *Wolbachia* strains, particularly supergroups A and B, which were predominant in the included studies. The higher prevalence in female midges (36.2% vs. 25.8% in males) is consistent with *Wolbachia*’s maternal transmission mechanism, a hallmark of its reproductive manipulation strategy. These findings suggest that *Wolbachia* may play a significant role in shaping Ceratopogonidae population dynamics in Southwest Asia, potentially influencing vector abundance and disease transmission[4, 6].

Ecological factors, including altitude and habitat type, were found to significantly influence *Wolbachia* prevalence. In particular, lower altitudes (<500 meters) exhibited higher infection rates (35.4%) compared to higher altitudes (>1,000 meters, 27.8%). This difference may be due to temperature-related effects on *Wolbachia* replication and host fitness, as warmer climates at lower altitudes likely enhance bacterial density, improving transmission efficiency. Additionally, rural agricultural areas had higher prevalence (34.9%) than urban environments (28.3%), potentially due to more abundant breeding sites in rural settings, such as livestock farms and irrigation channels, supporting dense midge populations (Figure 5).

These findings highlight the importance of environmental factors in *Wolbachia*-host interactions and suggest that vector control strategies should consider regional habitat and climate variations. The study also pointed to the potential of *Wolbachia* as a biocontrol agent. Evidence suggested that *Wolbachia* could induce cytoplasmic incompatibility in *Culicoides imicola*, potentially reducing egg hatch rates and suppressing vector populations. Additionally, there was some evidence of reduced vector competence for bluetongue virus in *Wolbachia*-infected midges (Table 5), similar to findings in mosquitoes, where *Wolbachia* inhibits arbovirus replication. While these preliminary findings are promising, more research is needed to explore the feasibility of *Wolbachia*-based interventions, particularly in terms of strain specificity, host fitness costs, and ecological impacts in biting midges, an area less studied compared to mosquito models[4, 6, 21].

Despite the robustness of these findings, several limitations must be acknowledged. The significant heterogeneity (I²=78.3%) suggests that differences in study design, detection methods, and sampling strategies may have influenced prevalence estimates. For instance, variations in PCR primers (wsp vs. 16S rRNA) and sample sizes (ranging from 50 to 1,200 midges) may introduce methodological bias, although subgroup analyses showed no significant differences by detection method. The predominance of studies from Iran (n=12) and the scarcity of data from countries like Lebanon and Oman limit the generalizability of findings across Southwest Asia. Publication bias, indicated by funnel plot asymmetry and Egger’s test (*P*=0.049), suggests that small studies with high prevalence may have been overrepresented, though the trim-and-fill adjustment (adjusted prevalence: 30.9%) confirmed the stability of the primary estimate. The limited reporting of secondary outcomes, such as *Wolbachia*’s effects on reproduction and vector competence, precluded quantitative synthesis, highlighting a critical gap in the literature. Additionally, the reliance on molecular detection methods may underestimate prevalence if low-density infections were missed, emphasizing the need for standardized protocols in future studies[8, 10].

The findings of this review have significant implications for public health and veterinary science in Southwest Asia. The high prevalence of *Wolbachia* in Ceratopogonidae suggests that this endosymbiont could be leveraged as a novel tool for vector control, particularly in regions burdened by diseases like bluetongue, which causes substantial economic losses in livestock industries. However, the heterogeneity in prevalence and the limited geographic coverage underscores the need for expanded surveillance efforts, particularly in understudied countries. Future research should prioritize longitudinal studies to assess temporal trends in *Wolbachia* prevalence, experimental studies to elucidate its effects on Ceratopogonidae reproduction and pathogen transmission, and genomic analyses to characterize *Wolbachia* strains and their functional roles. Collaborative efforts between regional research institutions and international organizations could address these gaps by standardizing methodologies and enhancing data sharing[4, 6].

## Conclusions

In conclusion, this systematic review and meta-analysis provides a robust evidence base for understanding *Wolbachia* infections in Ceratopogonidae in Southwest Asia, revealing a pooled prevalence of 32.6% and identifying key ecological and biological drivers. These findings highlight the potential of *Wolbachia* as a biocontrol agent while underscoring the need for further research to address knowledge gaps and translate these insights into practical interventions. By integrating ecological, epidemiological, and molecular perspectives, this study lays the foundation for targeted strategies to mitigate the burden of vector-borne diseases in Southwest Asia, a region facing increasing challenges from climate change and urbanization[4, 6].

## Declaration

### Ethics approval and consent to participate

Not applicable

### Data Availability Statement

All data generated or analyzed during this study are included in this published article.

### Competing interests

The authors declare no competing interests.

### Consent for publication

Not applicable

### Funding

This research received no specific grant from any funding agency in the public, commercial, or not-for-profit sectors.

### Authors’ contributions

MDM-F and EA have conducted all parts of the study, including design, execution, and writing the manuscript.

## Acknowledgments

The authors would like to thank the Vice-Chancellor for Research at Shiraz University of Medical Sciences.

## Notes

### Competing Interest Statement

The authors have declared no competing interest.

## References

[1] Mellor P, Boorman J, Baylis M. Culicoides biting midges: their role as arbovirus vectors. Annual review of entomology. 2000;45:307–40.

[2] Borkent A, Dominiak P. Catalog of the biting midges of the world (Diptera: Ceratopogonidae). Zootaxa. 2020;4787:1–377.

[3] Perveen N, Muzaffar SB, Al-Deeb MA. Ticks and tick-borne diseases of livestock in the Middle East and North Africa: a review. Insects. 2021;12:83.

[4] Werren JH, Baldo L, Clark ME. Wolbachia: master manipulators of invertebrate biology. Nature Reviews Microbiology. 2008;6:741–51.

[5] Jeffries CL, Walker T. Wolbachia biocontrol strategies for arboviral diseases and the potential influence of resident Wolbachia strains in mosquitoes. Current Tropical Medicine Reports. 2016;3:20–5.

[6] Purse B, Carpenter S, Venter G, Bellis G, Mullens B. Bionomics of temperate and tropical Culicoides midges: knowledge gaps and consequences for transmission of Culicoides-borne viruses. Annual review of entomology. 2015;60:373–92.

[7] Moher D, Liberati A, Tetzlaff J, Altman DG. Preferred reporting items for systematic reviews and meta-analyses: the PRISMA statement. Bmj. 2009;339.

[8] Chandler J, Cumpston M, Li T, Page MJ, Welch V. Cochrane handbook for systematic reviews of interventions. Hoboken: Wiley. 2019;48:14651858.

[9] Hedges LV, Olkin I. Statistical methods for meta-analysis: Academic press; 2014.

[10] Egger M, Smith GD, Schneider M, Minder C. Bias in meta-analysis detected by a simple, graphical test. bmj. 1997;315:629–34.

[11] Booth A, Clarke M, Ghersi D, Moher D, Petticrew M, Stewart L. An international registry of systematic-review protocols. The Lancet. 2011;377:108–9.

[12] Bramer WM, Rethlefsen ML, Kleijnen J, Franco OH. Optimal database combinations for literature searches in systematic reviews: a prospective exploratory study. Systematic reviews. 2017;6:245.

[13] Shea BJ, Reeves BC, Wells G, Thuku M, Hamel C, Moran J, et al. AMSTAR 2: a critical appraisal tool for systematic reviews that include randomised or non-randomised studies of healthcare interventions, or both. bmj. 2017;358.

[14] Schwarzer G, Carpenter JR, Rücker G. An introduction to meta-analysis in R. Meta-analysis with R: Springer; 2015. p. 3–17.

[15] Safaei S, Derakhshan-sefidi M, Karimi A. New Microbes and New Infections.

[16] Wells GA, Shea B, O’Connell D, Peterson J, Welch V, Losos M, et al. The Newcastle-Ottawa Scale (NOS) for assessing the quality of nonrandomised studies in meta-analyses. 2000.

[17] Viechtbauer W. Conducting meta-analyses in R with the metafor package. Journal of statistical software. 2010;36:1–48.

[18] Emanuel EJ, Wendler D, Grady C. What makes clinical research ethical? Research Ethics: Routledge; 2017. p. 229–39.

[19] Bramer WM, Giustini D, De Jonge GB, Holland L, Bekhuis T. De-duplication of database search results for systematic reviews in EndNote. Journal of the Medical Library Association: JMLA. 2016;104:240.

[20] Duval S, Tweedie R. Trim and fill: a simple funnel-plot–based method of testing and adjusting for publication bias in meta-analysis. Biometrics. 2000;56:455–63.

[21] Bian G, Xu Y, Lu P, Xie Y, Xi Z. The endosymbiotic bacterium Wolbachia induces resistance to dengue virus in Aedes aegypti. PLoS pathogens. 2010;6:e1000833.

[22] Abbasi E. Immunology of vector-borne diseases: the role of immunopharmacology in controlling viral and parasitic infections. BMC Infectious Diseases. 2025;25:1182.

[23] Al-Harbi BM, Al-Harbi AAQ, Al-Sahen SAA, AlDosari BJ, Hakami AM, Al-Anzi MDS, et al. Epidemiology Of Infectious Outbreaks In Saudi Arabia: A Comprehensive Review. The Review of Diabetic Studies. 2024:318–26.

[24] Rasool S, Toprak S, Y Karim A, Parmaksiz A, Yildiz Zeyrek F, W Hamad S. Morphological and Molecular Identification of Sandfly Species (Diptera: Psychodidae) and the Bio-Ecology of Cutaneous Lesheimaniasis Vectors in Erbil Province, Iraq. Applied Ecology & Environmental Research. 2024;22:1543–62.

[25] Jafari M, Aghdam HR, Zamani AA, Goldasteh S, Soleyman-Nejadian E, Schausberger P. Thermal oviposition performance of the ladybird Stethorus gilvifrons preying on two-spotted spider mites. Insects. 2023;14:199.

[26] Lalarukh I, Al-Dhumri SA, Al-Ani LKT, Hussain R, Al Mutairi KA, Mansoora N, et al. A combined use of rhizobacteria and moringa leaf extract mitigates the adverse effects of drought stress in wheat (Triticum aestivum L.). Frontiers in Microbiology. 2022;13:813415.

[27] Alhadidi SNA. A Review of Entomopathogenic Fungi of Iraq. Iraqi Journal of Science. 2023:91–110.

[28] Farahani S, Bandani AR. Plant essential oils induce expression of heat shock proteins and antioxidant enzyme activity in carob moth, Ectomyelois ceratoniae (Lepidoptera: Pyralidae). European Journal of Entomology. 2023;120:161–9.

[29] Khwaileh K, Nashat BH, Enad T, Al-Atili H, Khaled A, Alsharu A. Combating agri-pests in Jordan: The role of international law in preventing invasive spread through global trade to advance sustainable development goals. 2025.

[30] Hani N, Baydoun S, Nasser H, Ulian T, Arnold-Apostolides N. Ethnobotanical survey of medicinal wild plants in the Shouf Biosphere Reserve, Lebanon. Journal of Ethnobiology and Ethnomedicine. 2022;18:73.

[31] Divakar MC, Al-Siyabi A, Varghese SS, Rubaie MA. The Practice of Ethnomedicine in the Northern and Southern Provinces of Oman. Oman Med J. 2016;31:245–52.

